# Development and analytical validation of an enzyme-linked immunosorbent assay based on baculovirus recombinant LipL32 protein antigen for the accurate detection of canine leptospirosis

**DOI:** 10.1101/358861

**Authors:** Carolina Orozco-Cabrera, Gilberto López-Valencia, Sergio Arturo Cueto-González, José Guadalupe Guerrero-Velázquez, Kattya Moreno-Torres, Kelvin Orlando Espinoza-Blandón, Nohemí Castro-Del Campo, Soila Maribel Gaxiola-Camacho, Sergio Daniel Gómez-Gómez, Enrique Trasviña-Muñoz, Cinthya Torres-Guzmán, Francisco Javier Monge-Navarro

## Abstract

*Leptospira* infects a wide range of companion, domestic and wild animal species, shedding the spirochetes into the environment via urine. Dogs become infected by direct or indirect contact with wild or domestic infected animal reservoirs increasing the risk of zoonotic transmission of the disease. The microscopic agglutination test has been used as the gold standard for the diagnosis of leptospirosis but has low sensitivity and is technically complex. Several ELISA tests have been developed based on recombinant proteins of *Leptospira* for the diagnosis of leptospirosis with similar or higher specificity and sensitivity levels than the microscopic agglutination test. Here, we developed and analytically validated an ELISA test based on recombinant LipL32 protein of *Leptospira* expressed in baculovirus. The LipL32 protein was successfully adapted in an indirect ELISA using dog plasma samples. Optimization of the ELISA resulted in a P/N ratio of 7.18 using only 5 ng of rLipL32 per well. Inter-assay and intra-assay variation showed a CV of 3.96% and 6.98% respectively, suggesting that the ELISA-LipL32 is highly reproducible. When tested with field samples, concordance of the ELISA-LipL32 with a real-time PCR, positive concordance was 100%. Our results indicate that the ELISA-LipL32 has the potential to be used by veterinarians and public health investigators as a safe, rapid, inexpensive and reliable method for the early diagnosis of *Leptospira* infection in dogs. Additional studies are still required for clinical validation on field samples under different epidemiological scenarios.

## Introduction

Leptospirosis is the most widespread zoonotic disease in the world affecting over 1 million people every year with a case fatality between 5 and 10% of cases. The majority of the reports for morbidity and mortality of leptospirosis occur in adults ranging 20-49 years of age from poor regions of the world and from regions lacking of appropriate surveillance programs for this disease (3). In Latin America, leptospirosis are routinely reported in 24 out of 48 countries but only 18 countries from that region had a government mandatory notification system, resulting in a limited geographical knowledge of the epidemiology of leptospirosis and an underestimation of the real morbidity and mortality of the disease in the Americas (4). Leptospirosis is caused by pathogenic bacteria of the genus *Leptospira*. Genotyping classification recognizes at least 20 species, including 9 pathogenic, 5 intermediate pathogenic and 6 saprophytic species of *Leptospira* (17). Pathogenic and intermediate pathogenic leptospiras chronically infect a wide range of companion, domestic and wild animal species, shedding the spirochetes into the environment via urine, where they can survive for weeks or months in water or humid soil until an incidental host got infected upon contact and penetration through mucosal membranes, skin abrasions or lacerations. Humans can become accidentally infected by contact with contaminated soil or water and from urine of infected animal reservoirs (13). Leptospirosis produced by *Leptospira interrogans* serovars *Icterohaemorrhagiae* and *Canicola* which are considered the main serovars affecting dogs (17). Dogs are frequently exposed to zoonotic pathogens and become infected by direct or indirect contact with other wild or domestic infected animal reservoirs in their immediate outdoor environment. This contamination might lead to an infection that bring *Leptospira* into the household, increasing the risk of zoonotic transmission of the disease between dogs and their owners by direct contact with leptospiras shed in the urine or present in another common contaminated source like water (8). Early detection and diagnosis of *Leptospira* infection must be achieved in order to establish an appropriate antibiotic therapy. For many years, the microscopic agglutination test (MAT) has been used as the gold standard for the diagnosis of leptospirosis; however, given its technical complexity, time consuming and the biosecurity risk that represents to the personnel handling pure strains of pathogenic *Leptospira* to obtain whole cell antigens, MAT is performed only at reference laboratories where highly trained personal and biosecurity measurements are available (6, 16). It has been demonstrated that MAT has low sensitivity when is used within the first 10 days of infection and require the testing of acute and convalescent samples to obtain better sensitivity levels, limiting the use of MAT only for confirmation of infection with *Leptospira* once the antibodies have raised to detectable levels in the patient (12). To overcome the limitations of MAT, several ELISA platforms for the diagnosis of leptospirosis has been developed based on the use of whole cell antigens for detection of antibodies (10, 16), or by the use of specific antibodies to detect circulating leptospiras in serum or plasma samples (5, 10). More recently, several ELISA platforms have been developed based on highly specific recombinant proteins of *Leptospira* for the diagnosis of leptospirosis in both humans (5, 6, 10), and different animal species such as bovine (2), equine (19), swine (9), and dogs (20), with similar or higher specificity and sensitivity levels than MAT. The use of the adhesion protein LipL32, a highly conserved immunodominant protein expressed only in pathogenic species of *Leptospira* (14), has been successfully adapted into different serological platforms for the diagnosis of leptospirosis in humans (6) and animals (2, 9, 20). In this study, we report the development and analytical validation of an ELISA system based on the LipL32 protein of *Leptospira* expressed in a baculovirus expression system. The recombinant LipL32 protein (rLipL32) was abundantly produced in insect cell culture, purified using Ni-NTA affinity chromatography and successfully adapted to be used as solid phase antigen for development of an indirect ELISA for the accurate detection of canine leptospirosis in dog plasma samples.

## Materials and methods

### Location of the study

The standardization and analytical validation of the ELISA-rLipL32 system were performed at the Institute for Research in Veterinary Sciences from the Autonomous University of Baja California, campus Mexicali, in Northwest Mexico. Mexicali is the capital city of the state Baja California (32°39’48’’N 115°28’04’’W) and shares the international border line with the state of California in the southwest United States of America.

### Study population

A total of 57 plasma samples from stray dogs captured by the Mexicali County Animal Control Center (MCACC) were included in this work. This group of samples was used in a previous study for the development and analytical validation of a *Leptospira* LipL32 real-time PCR (RT-PCR-LipL32) diagnostic platform (data not published).

### Recombinant protein LipL32

The design of rLipL32 protein was based on the sequence of *Leptospira interrogans* strain 200901482 LipL32 (GenBank: JN683904.1), a 474 base pair (bp) gene that encodes for a protein of 158 amino acids (GenBank: AEZ53279.1). Expression and synthesis of rLipL32 was performed by GenScript (Piscataway, NJ, USA) using a baculovirus expression system. Briefly, the original sequence JN683904.1 was codon optimized and an 8-His tag added at the amino terminal of the DNA molecule. The optimized target DNA sequence of LipL32 was synthesized and sub-cloned into the pFastBac1 vector (Invitrogen, Carlsbad CA, USA) for insect cell expression. Max Efficiency DH10Bac competent cells were used for the generation of recombinant bacmid DNA and transfected into Sf9 insect cell culture. Sf9 cell cultures were incubated in Sf-900 II serum-free, protein-free insect cell culture medium for 5 to 7 days at 27 °C. Cell cultures were harvested and centrifuged and the collected supernatant was designated as the P1 viral stock. A P2 amplified viral stock was produced for infection. For expression of rLipL32, the P2 amplified viral stock was optimized for infection of Sf9 cell cultures at 0.5, 3, and 5 moieties of infection (MOI) and harvested at 48, 72, and 96 hours post infection. Cell pellets were harvested and lysed using a proper cell lysis buffer. Cell lysates were incubated with Ni-NTA columns to capture rLipL32 target protein. Eluted fractions were pooled, dialyzed and analyzed by SDS-PAGE and western blot using standard protocols for molecular weight and purity measurements. The primary antibody for western blot was a mouse ‐anti ‐His horse radish peroxidase (HRP) labeled monoclonal antibody. Protein concentration was determined by the Bradford protein assay using bovine serum albumin (BSA) as a standard. A batch of 3.6 milligrams (mg) of rLipL32 with a concentration of 0.20 mg/milliliter and purity of 80% estimated by densitometry analysis of the Coomassie blue-stained SDS-PAGE was produced and delivered by GenScript.

### Positive and negative controls

Positive controls were obtained from a pool of six plasma fractions of whole blood samples collected from stray dogs that resulted positive to a RT-PCR-LipL32 developed and analytically validated in our lab in a previous study (data not shown). These plasma samples were selected as positive controls because in the real-time PCR they showed an early cycle of amplification (Threshold cycle <20) and developed the highest relative fluorescence unit (RFU) score in their respective real-time PCR test run. Since antibodies to *Leptospira* are produced between the 6th to the 10^th^ day after, reaching peak levels at approximately 4 weeks after infection (12, 14) thus, a positive PCR could be indicative of current infection with presence of detectable circulating antibodies to LipL32. A normal canine serum commercially available (Catalog number S1757, Sigma-Aldrich Corp. St. Louis, MO, USA) was used as negative control.

### rLipL32 ELISA protocol

The rLipL32 protein was standardized for use as solid phase antigen for the detection of antibodies in an indirect ELISA system. Optimal antigen concentration, serum samples, and enzyme conjugate antibody dilutions, were determined through standard checkerboard titration procedures (15). Briefly, rLipL32 protein was diluted in carbonate-bicarbonate buffer pH 9.6 at a concentration of 200, 100, 50, 25, 10.0 and 5.0 ng of rLipL32 protein antigen per well. After incubation overnight at 4°C plates were ready to use. To run the tests, plates were washed five times with phosphate buffered saline solution pH 7.5 containing 0.05% Tween 20 (PBST). Positive and negative controls were diluted two-fold from 1:25 to 1:200 and added in duplicates to appropriate wells. After incubation at 37°C for 1 hour, plates were washed as described before and 100 µl of anti-dog IgG HRP conjugate diluted 1:12,500, 1:25,000, 1:50,000 and 1:100,000 in PBTS were added to individual plates. After another round of incubation and washing, 100 µl of substrate solution (TMBS) (Sigma-Aldrich Corp., St. Louis, MO, USA) were added to each well and color development stopped with 30 µl per well of 0.16M sulfuric acid solution after 10 minutes of incubation at room temperature (RT). Optical density values were obtained at 450 nm using an automated ELISA reader. The selection criteria for the best performance of the ELISA system was determined based on the amount of antigen per well, use of blocking solution, dilution of serum and dilution of the enzyme conjugate that resulted in the maximum amplitude of difference between the mean OD value of the positive control serum over the mean OD value of the negative control serum, expressed as the positive/negative ratio (P/N ratio).

### Precision of the ELISA-rLipL32

Ten plasma samples positive to the RT-PCR-LipL32 and 10 aliquots of the canine sera previously used as negative control in the ELISA standardization protocol were tested in duplicates at random positions across the 96 wells of the microtiter plates during six non-consecutive days. From the OD obtained, standard deviation (SD) and coefficient of variation (CV) were calculated to obtain the repeatability of plasma samples tested on the same plate (intra-assay variation) and on different plates at different times (inter-assay repeatability). A CV value less than 20% was considered to produce acceptable variation (1).

### Dynamic range of the ELISA-LipL32

Lower and upper limit of detection and dilution linearity were calculated to establish the working range of the ELISA-LipL32 that produce a reliable result. Two-fold serial dilutions of the plasma positive control pool were tested from dilution 1:25 (working dilution) to 1:6,400. One set of the canine negative control serum diluted 1:25 and one set of blanks of PBST were included in the test run. All positive, negative and blank samples were run in octuplicates in a microtiter plate.

### ELISA-rLipL32 with field samples

A group of 57 plasma samples from stray dogs captured by the County Animal Control Center (MCACC) was tested to detect antibodies to *Leptospira* using the optimized ELISA-rLipL32. Total concordance rate (TC), positive concordance rate (PC) and negative concordance rate (NC) were calculated and compared with the results of the RT-PCR-LipL32 method using the following formulas: (TC) = (TP + TN)/total number of serum samples tested x 100; Positive concordance rate PC (%) = (TP/total number of positives in the RT-PCR-LipL32) x 100, where TP is the number of positives in both tests, and Negative concordance rate NC (%) = (TN/total number of negatives in the RT-PCR-LipL32) x 100, where TN is the number of negatives in both tests.

### Data analysis

Data analysis was performed using the Statistical Analysis Systems (SAS) version 9.4.

## Results

### ELISA-rLipL32 protocol optimization

The maximum P/N ratio with the lowest background noise of the ELISA-LipL32 was established in 7.18 with a mean OD of 1.487 for the positive control samples and a mean OD of 0.207 for the negative control (Table 1). These results were obtained using 5 ng of rLipL32 protein antigen per well, the serum samples diluted 1:25 and the conjugate diluted 1:50,000. Incubation of primary and secondary antibody was performed at 37°C for 1 hour each, followed by 10 minutes of incubation with TMBS at RT.

**Table 1.**
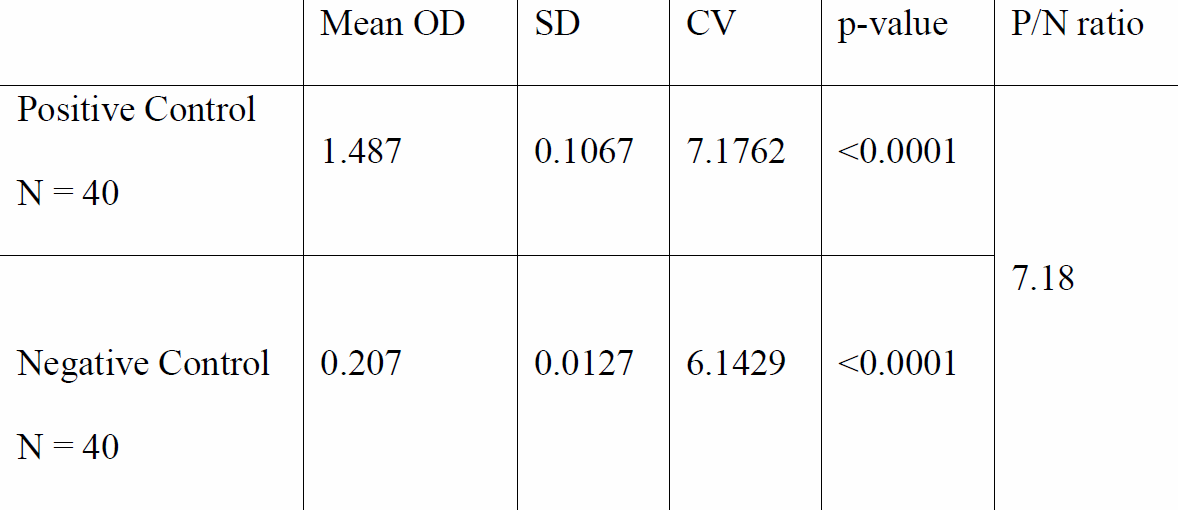
Optimization of the ELISA-LipL32. The amplitude between the OD of positive and negative samples showed a good optimization of the reagents, demonstrating the capacity of the test to produce a clear differentiation between the positive from the negative control samples with minimal background. Mean OD = Mean optical density SD = Standard deviation CV = Coefficient of variation P/N ratio= Positive / negative ratio

### Precision of the ELISA-LipL32

Ten positive plasma samples and ten negative aliquots of canine sera were tested during six nonconsecutive days for intra-assay and inter-assay variation to evaluate the repeatability of the ELISA. Results showed an intra-assay CV of 6.98% (4.13% − 11.20%) and an inter-assay CV of 3.96% (2.12% − 8.60%). These data suggests that ELISA-LipL32 has the capacity to produce reproducible results with low variation (Table 1).

### Dynamic range of the ELISA-LipL32

The higher OD detected during intra-assay and inter-assay experiments was 1.870, which corresponds to a positive pooled plasma sample diluted 1:25 and it was established as the upper limit of detection. The lower OD detected was 0.189, which corresponds to a positive pooled plasma sample diluted 1:6,200 and it was regarded as the lower limit of detection (Table 3).

### Dilution linearity

To obtain the dilution linearity of the ELISA-LipL32, the mean OD of two-fold sample dilution series were plotted and a regression curve (R^2^) calculated. The results shown a linearity with an R^2^ value of 0.9901 (p<0.001), demonstrating that rLipL32 is capable to allow the reliable determination of antibody levels in diluted samples within the OD dynamic range established for this method (Table 3).

### Concordance of the ELISA-LipL32 with the RT-PCR-LipL32

A total of 57 plasma samples previously tested using the RT-PCR-LipL32 method were also tested with the ELISA-LipL32 to determine the level of concordance between both tests. The results of the ELISA showed a total concordance rate of 77.19 %, a positive concordance rate of 100.0 % and 0 % of negative concordance rate (Table 2).

**Table 2.** Concordance of the RT-PCR-LipL32 and the ELISA-LipL32. Plasma samples from stray dogs (N= 57) tested positive to LipL32 by RT-PCR were compared with the results of the ELISALipL32 and concordance rates calculated showing a PC rate of 100%. RT-PCR can amplify *Leptospira* DNA from blood for a short (10 days) period of time. Antibodies become detectable during the first week of infection and remain detectable for weeks or months, making the ELISA identify to detect more positive samples.

|  | Positive | Negative | Total concordance |
| --- | --- | --- | --- |
| RT-PCR-LipL32 | 45/57 (78%) | 12/57 (21%) | 77.19% |
| ELISA-LipL32 | 57/57 (100%) | 0/57 (0%) |  |
| Concordance | 100.0% | 0.0% |  |

**Table 3.**
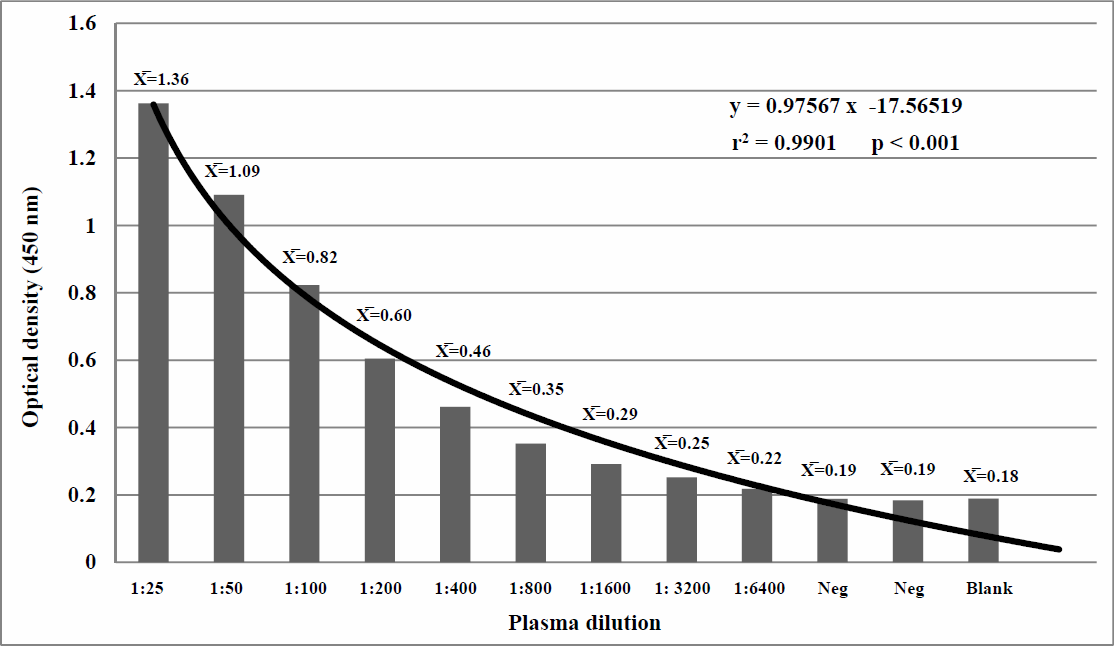
Dilution linearity of the ELISA-LipL32. Two-fold serial diluted plasma samples were tested using the optimized ELISA-LipL32. Results showed a strong linearity (R^2^ =0.9901) in diluted samples with no effect over the precision of the test. The mean OD at dilution 1:6400 is just above the mean OD of the negative control and blank samples.

## Discussion

Accurate detection and diagnosis of a*Leptospira* infection is important to the prompt initiation of the antibiotic therapy and to minimize the spread of the disease. A delay in the diagnosis represents a significant challenge for human and veterinary public health authorities facing the possibility of an epidemic outbreak of leptospirosis, especially in regions where the pathogen is emergent. MAT is considered the reference serological method used to diagnose leptospirosis. However, because of the low sensitivity of the test MAT is considered a poor gold standard for comparison of other diagnostic tests for leptospirosis (12). Also, this procedure is laborious, expensive, requires of stocks of live leptospiras and must be carried out in specialized reference laboratories with high level bio-containment facilities; consequently it is rarely used in most developing countries (6). PCR based molecular methods are also very sensitive and specific for the diagnosis of *Leptospira* but, given the short period of the bacteriuria in infected dogs, the usefulness of this technology for early diagnosis of the disease is limited to the first 10-12 days after infection (11). Also, RT-PCR diagnosis of *Leptospira* requires well equipped laboratories, which might be beyond the resources and capabilities of clinical and diagnostic laboratories in many developing countries. Antibody detection by ELISA for the diagnosis of *Leptospira* is widely used. The emergence of leptospirosis world-wide resulted in an increased demand for safe, standardized and validated diagnostic methods for the diagnosis of leptospirosis (20) (14). The development of recombinant proteins is one approach that is suitable for the safe production of high quality ELISA reagents given that are easily expressed, produced and standardize with high degree of consistency between batches without the biosecurity risks associated extraction of leptospiral antigen from pure microbiological culture preparations (6, 20). Here, we developed an ELISA based on a baculovirus recombinant LipL32 protein with potential application as diagnostic antigen in an indirect ELISA system for detection of antibodies in dogs plasma samples. The LipL32 protein was chosen as an antigen because it has been shown to be an immunodominant protein exclusively expressed for pathogenic species of *Leptospira*. The cloning, expression and purification methods for the production of the rLipL32 protein described in this work resulted in a technically simple, cost effective production of a highly reliable, very stable and noninfectious antigen for ELISA. The rLipL32 protein strongly bound to the ELISA plates generating minimal background, thus producing an effective differentiation between the positive from the negative controls tested with a P/N ratio of 7.18. Considering the working dilution of the rLipL32 protein antigen for the ELISA optimized at 5 ng per well, a batch of 1 milligram of and 80% purity rLipL32 protein like the one tested here, would be enough to test approximately 100,000 duplicate plasma or serum samples for detection of antibodies to *Leptospira,* making the ELISA-LipL32 an inexpensive platform to the accurate detection of leptospirosis in dogs and possibly in other animal species. Dilution linearity experiments resulted in a strong correlation (R2=0.9901; p<0.001), determining to which extent the response of the diluted sample is linear through the working range of the test with no effect over the precision (1). This linearity strength will be of great value for the analysis and characterization of the antibody response from animals living under different epidemiological situations such as urban or rural setting, stray dog or household dog, age, gender, vaccination status, antibiotic treatments and others variables that might help to achieve field validation of the ELISA-LipL32 in the future. Stray dogs are probably involved in the circulation of pathogenic *Leptospira* and could be playing a significant role in the transmission of leptospirosis acting as a vector from wild or feral reservoirs infecting dogs and humans in the household environment (8). Concordance rate experiments showed that the ELISALipL32 was capable to detect more (57/57) samples positive for antibodies to rLipL32 that the RTPCR (45/57) reference method. This can be explained because of the fact that after infection, *Leptospira* rapidly proliferates in the blood stream producing an acute phase that lasts for about 10 days when molecular diagnosis is capable to produce a positive result (20). On the other hand, antibodies can be detected in blood by the 6^th^ to 10th day post infection, which coincides with the clearance of *Leptospira* from the blood stream. Antibodies will continue to rise and reach peak levels 3-4 months after infection thus, more cases of leptospirosis can be detected by using serology than molecular methods (18). The high rate (100%) of plasma samples tested positive with the ELISA-LipL32 showing OD >1.00, strongly suggests evidence of active infection in the stray dog population and may pose a public health problem to humans (7). So far, the results of the present work indicate that the ELISA-LipL32 may be a reliable tool for early diagnosis of leptospirosis and has the potential to be used by veterinarians and public health investigators as a safe, rapid, inexpensive and reliable method for the early diagnosis of *Leptospira* infection in dogs, to study the epidemiology of this disease and to evaluate current sanitary strategies for control and prevention of leptospirosis. Further studies oriented to achieve clinical validation on field samples under different epidemiological scenarios are still required.

## Acknowledgments

This study was partially supported with funds from the 18va. Convocatoria Interna de Apoyo a Proyectos de Investigación from Universidad Autónoma de Baja California.

